# An integrated genomic framework for *Aeromonas* genomic species delineation using average nucleotide identify, core-genome phylogeny and digital DNA-DNA hybridization

**DOI:** 10.64898/2026.02.04.703925

**Authors:** Alex Chen Lu, Ruochen Wu, Ruiting Lan, Li Zhang

## Abstract

*Aeromonas* taxonomy has long been complicated by overlapping phenotypic, biochemical, and protein profiles. Here, we establish a robust genome-based framework for *Aeromonas* genomic species delineation. We analysed average nucleotide identity (ANI) across 4,366 available *Aeromonas* genomes and demonstrated that at 96% ANI threshold, skANI and fastANI generated too many clusters (65 and 57 respectively) and these clusters were not supported by core genome phylogeny. We identified 95.4% skANI (equivalent to 95.6% fastANI) as an operational threshold for the delineating *Aeromonas* genomic species. Using the 95.4% skANI threshold, we identified 44 ANI clusters among the 4,366 genomes, of which 43 clusters were genomic species supported by the core-genome phylogeny. Thirty-four of the 43 genomic species corresponded to existing taxonomic species, whilst the remaining nine are currently not recognised as taxonomic species. All recognised taxonomic species represented in the dataset retained their existing species designation except *Aeromonas mytilicola*, which was not separated from *Aeromonas rivipollensis* in both ANI clusters and the core-genome phylogeny. The digital DNA–DNA hybridisation (dDDH) values between the genomic species were below 70%, further supporting genomic species delineation. We further developed AeromonasGStyper, a genomic species typing tool that assigns query genomes based on ANI similarity to medoid genomes. In conclusion, this study establishes a genomic species framework for genome-based classification of *Aeromonas* and provides a practical approach for future genomic surveillance.

**Impact Statement:** *Aeromonas* species have gained increased attention as emerging human enteric pathogens. *Aeromonas* taxonomy has long been complicated by overlapping phenotypic, biochemical, and protein profiles. Although a 96% average nucleotide identity (ANI) threshold was proposed previously for *Aeromonas* species delineation, analysis of 4,366 *Aeromonas* genomes demonstrated that this threshold generated excessive genomic clusters that were not supported by the core-genome phylogeny. We identified 95.4% skANI (equivalent to 95.6% fastANI) as an operational threshold for delineating *Aeromonas* genomic species, supported by core-genome phylogeny and digital DNA–DNA hybridisation (dDDH). The framework identified 43 genomic species, including 34 corresponding to recognised taxonomic species and nine genomic species that do not correspond to currently recognised taxonomic species. In addition, our data showed that *Aeromonas mytilicola* was not separated from *Aeromonas rivipollensis* by ANI clustering and core-genome phylogeny, supporting further taxonomic reassessment of the distinction between these two species.

## Introduction

The genus *Aeromonas* are a highly diverse group of bacteria that occupy a wide range of aquatic, environmental and host-associated niches (1). Early research recognised *Aeromonas hydrophila* as a highly virulent opportunistic pathogen responsible for severe disease outbreaks in fish, resulting in substantial economic losses in global aquaculture (2). Several other *Aeromonas* species have since been identified as causative agents of disease outbreaks in fish and other farmed animals, highlighting their ecological diversity and pathogenic breadth of this genus (3, 4). Beyond their importance in animal health, *Aeromonas* species are recognised as human pathogens. Some species are well-documented causes of human soft tissue, wound, and bloodstream infections, and their role in human gastrointestinal diseases is increasingly recognised (5, 6). Human infections are often associated with contaminated water, food or aquatic animals and can be especially severe in immunocompromised individuals, highlighting the public health relevance of the genus (5, 7, 8).

As *Aeromonas* species have increased in importance in relation to global aquaculture and public health, both molecular and genomic techniques have been used to identify and analyse species. However, there is a lack of clarity between existing taxonomy, literature and different genome-based identifications of *Aeromonas*. This is partially due to complications and limitations in classical identification methods. The 16S rRNA gene sequencing is unable to differentiate closely related *Aeromonas* species due to high conservation and interspecies recombination in the genus (9, 10). Similarly, phenotypic and biochemical identification is difficult due to overlapping characteristics between species (11). With the advent of next-generation sequencing, genome-based comparisons are increasingly used for *Aeromonas* species identification (12, 13).

For prokaryotic species delineation, genome-based pairwise comparative methods include digital DNA-DNA hybridisation (dDDH) and average nucleotide identity (ANI). dDDH is an *in silico* implementation of the traditional DNA-DNA hybridisation (DDH), which was considered a gold standard for species delineation, whereas ANI provides a genome-wide measure of nucleotide similarity between genomes (14). Both methods offer computational efficiency and the quantitative criteria for species classification (15–17). In bacteria, ANI thresholds of 95% to 96% are used for bacterial delineation, a threshold that corresponds to the DDH and dDDH cutoff of 70% for species delineation (18–20). In contrast, core genome phylogeny aims to evaluate the evolutionary phylogeny on the genus. However, the technique is much more computationally and time intensive, limiting its use in practical species identifications (18, 21).

Currently, k-mer based ANI methods are among the most widely utilised methods of genome-based species identification and are extensively applied in literature and in large-scale genome classification frameworks, including the National Centre of Biotechnology Information (NCBI) taxonomy and Genome Taxonomy Database (GTDB). However, NCBI and GTDB apply different ANI thresholds for species delineation, with GTDB and NCBI opting for threshold of 95% and 96% ANI respectively (20–22)(22, 23). K-mer based ANI approaches are much more efficient than alignment-based ANI methods, albeit at the cost of accuracy in comparison to evolutionary phylogeny (24, 25).

Studies evaluating ANI in comparison to genome-scale phylogeny at the genus level, dDDH, and existing *Aeromonas* taxonomy are limited. Colston et al. in 2014 examined 58 *Aeromonas* genomes and recommended 96% ANI cutoff as a practical threshold for the genus of *Aeromonas* (12). The number of *Aeromonas* genomes has increased greatly in recent years and multiple *Aeromonas* species have been proposed based on the 96% ANI and the 70% dDDH threshold (26–29). In addition, phylogenetic complexities have been observed within and between species such as the *Aeromonas media* complex, *Aeromonas sobria*, and *Aeromonas veronii*, which warrant analysis at a genus level (30, 31). It is essential to reassess how well the ANI-based species delineations correspond with genus-phylogeny and existing species definitions (27–29).

A stable genome-based species framework is not only a fundamental requirement for resolving *Aeromonas* taxonomy, but also critical for accurate disease surveillance (26). Precise species-level identification is necessary because distinct *Aeromonas* species may differ substantially in virulence factors, ecological niches, and pathogenic potential, with direct implications for risk assessment, and outbreak investigation (32–35).

In this study, we analysed how different ANI thresholds from ANI k-mer based tools and databases delineate species within the *Aeromonas* genus in comparison to core genome phylogeny. In addition, we compared the different species level delineations of *Aeromonas* with existing *Aeromonas* taxonomy. The dDDH values between species were also compared. Furthermore, we developed a tool that accurately assigns query genomes to genomic species. This study establishes a framework that supports future genome-based identification and pathogen surveillance within the *Aeromonas* genus.

## Materials and Methods

### *Aeromonas* genome retrieval and establishment of analysis datasets

Assembled *Aeromonas* genomes were downloaded from the NCBI database (https://www.ncbi.nlm.nih.gov/) (36). Additionally, publicly available paired end reads of *Aeromonas* were retrieved from the European Nucleotide Archive. Metadata including species assignment provided by genome submitters and the species stated by NCBI were also retrieved for taxonomic assignment. The short-read sequences were assembled using the Shovill v1.1.0 pipeline (https://github.com/tseemann/shovill). The program processed the raw reads using Trimmomatic, Lighter, Fast Adjustment of Short reads (Flash) and assembled them using strategic k-mer extension for scrupulous assemblies (SKESA) (37–40). The quality of all genomes was then assessed for outlying assembly metrics in total assembly length, contig number, N50 and the checkM metrics of completeness and contamination. The outlying ranges were calculated using interquartile ranges and are in Supplementary Table 1 (41).

### ANI-based genome comparison and clustering using average linkage clustering

ANI values were generated using the two k-mer based ANI tools fastANI v1.33 and skANI v0.3.2 respectively (15, 17). The pairwise results were compiled into dissimilarity matrices and clustered using the average linkage clustering algorithm from SciPy hierarchical clustering package (42). Clusters formed at ANI cutoffs ranging from 94% to 96.5%, with increments of 0.1%, were first examined and compared against each other. Graphs were then generated using pyplot from matplotlib v3.10.3 (43).

### *Aeromonas* genus core gene identification

A core gene database at the genus level for *Aeromonas* was constructed to perform further analysis.

A total of 393 genomes were selected to represent genus diversity in identifying the genus core genome. The 393 genomes comprised of 302 complete genomes, 78 chromosome level genomes, and 13 draft genomes for species without complete genomes. PROKKA was used to locate open-reading frames from the core genome dataset, from which the resultant GFF files were input into Roary v3.13 to define the *Aeromonas* genus core loci (44, 45). Roary v3.13 was run at multiple BLASTp percentage identities and gene presence ranges to identify potential settings for identifying core genes. The presence of core loci in each genome was determined by identifying genes with ≥ 85% minimum BLASTp percentage identity and 100% gene presence in the 393 genomes. Core genes that were paralogous or orthologous but were split into multiple fragments by erroneous assembly and/or annotation were identified and subsequently removed using a previously described Python script. The core genes and allelic variations were then combined into a core gene database (46, 47).

### Phylogenetic analysis based on genus core genome alignments

Core genome phylogenetic trees were then constructed based on the genus level core genome identified. Allele sequences of the *Aeromonas* genus core genes were called from all genomes using the core gene database. The individual alleles were then constructed into core genomes sequences by concatenating alignments of individual core loci. To reduce computational complexity, representative sets of genomes were used for phylogenetic analyses. The subsampling involved representing larger species clusters with a maximum of 300 genomes and keeping all genomes from species clusters with less genomes. Phylogenetic trees based on the core genome sequences were constructed using IQTREE v3.0.1 (48). Branch supports were assessed with non-parametric bootstraps at 1,000 replicates. The resulting phylogenetic trees were visualised using GrapeTree v1.5.0 (49).

### Species Identification of *Aeromonas* genomes

Multiple different methods were used to assign species identities to the *Aeromonas* genomes. The species identification prescribed by NCBI taxonomy and GTDB database were considered alongside species level taxonomic descriptions listed in published literature. Discrepancies between species identifications were evaluated in relation to the core genome phylogeny branches and ANI clusters generated using different ANI thresholds.

### Medoid genome identification

The pairwise dissimilarity matrix of ANI values and the clusters formed at the different ANI cutoffs were used as input for the Partition Around Medoids (PAM) algorithm. The FasterPAM algorithm in the package Fast k-medoids clustering in Python at default setting was used to identify the medoid genomes of each ANI cluster (50). For each cluster, the medoid genome was identified as the genome with the lowest dissimilarity score to all other members of the cluster.

### dDDH analysis

dDDH was also analysed in relation to species boundaries, with dDDH values calculated through the GGDC web service tool with the recommended values of Formula 2 taken as output (16). For interspecies comparisons, one medoid genome representing each genomic species were selected, and pairwise dDDH values were computed between medoid genomes. These comparisons were used to assess separation between genomic species. To evaluate the reliability of the 70% dDDH threshold for species delineation in the genus *Aeromonas*, dDDH values of representative strains within the same species were also examined.

### Development of an *Aeromonas* genomic species typing tool

The tool *Aeromonas* Genomic Species typer (AeromonasGStyper) was developed to assign *Aeromonas* species to query genomes based on their ANI similarity to medoid genomes found at the 95.4% skANI and 95.6% fastANI cutoff. The python program can utilise fastANI or skANI and can be found at (https://github.com/lizhanglab/AeromonasGStyper) (15, 17).

## Results

### Aeromonas genomes

*Aeromonas* genomes released before May 2026 in the NCBI and ENA databases were retrieved. The 4,366 genomes that have passed quality checks were used for analysis in this study. See methods and Supplementary Table 1 for quality metrics used and Supplementary Table 2 for the full list of genomes.

### Comparison of *Aeromonas* genome ANI clusters generated by fastANI and skANI tools

Pairwise ANI values between *Aeromonas* genomes were generated using both fastANI and skANI. The genomes were then clustered at ANI cutoff increments of 0.1% using average linkage clustering. Comparison between the tools showed that skANI thresholds produced clusters comparable to fastANI clusters when the fastANI threshold was reduced by 0.2% (Figure 1). Specifically, skANI thresholds of 94.6-94.9% and 95.1-95.4% generated clusters identical to those obtained using fastANI thresholds of 94.8-95.1% and 95.3%-95.6%, respectively.

**Figure 1.**
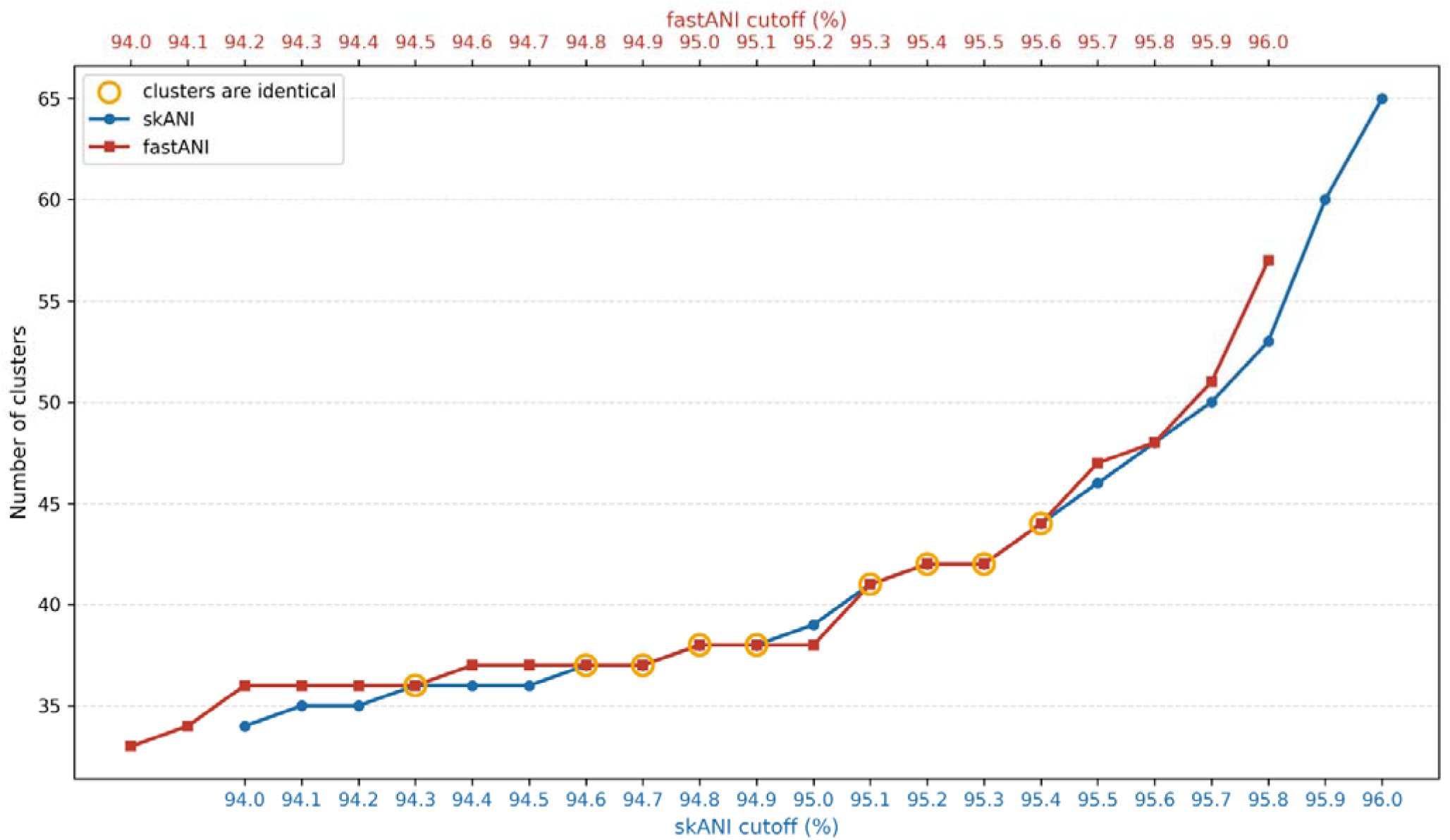
Number of genome clusters delineated from 4,366 genomes at different average nucleotide identity (ANI) cutoffs generated using skANI and fastANI. ANI clusters were generated by average-linkage clustering of 4,366 *Aeromonas* genomes using fastANI and skANI. The clusters between 94.6% to 95.4% skANI cutoffs match for most values between 94.8% to 95.6% fastANI cutoffs. The x-axis shows ANI cutoffs ranging from 94% to 96% for skANI (blue), with a 0.2% offset for fastANI (red). The y-axis shows the number of ANI clusters. Yellow circles indicate threshold pairs where both the number of clusters and the genomes assigned to each cluster are identical between the two methods.

The threshold of 96% ANI was previously proposed for species delineation in the *Aeromonas* genus (12). However, at this threshold, both fastANI and skANI formed excessive clusters, with 57 and 65 clusters respectively (Figure 1, Supplementary Figure 1). Other points in the 95-96% ANI threshold used for *Aeromonas* species delineation were considered instead.

No ANI threshold delineated all taxonomic species present in the dataset, but the four skANI cutoffs from 95.1-95.4% produced identical fastANI clusters more closely aligned with existing taxonomy.

To confirm whether the ANI clusters at different skANI and fastANI values were phylogenetically valid, and which threshold was most suitable for species delineation, comparison of ANI clusters with the core genome phylogeny was required.

### Comparison of core genome phylogeny with different ANI thresholds

For the construction of the core genome phylogenetic tree, a core genome first had to be established. To account for diversity in the genus, the core genome was determined using a total of 393 genomes representing 33 *Aeromonas* species recognised in NCBI taxonomy. A total of 705 genes initially passed core gene selection using Roary. Subsequently, 28 paralogous genes and 4 genes with overlapping regions were identified and removed. A final *Aeromonas* genus core genome comprising 673 genes and corresponding alleles from the 393 genomes was established.

The core genome phylogenetic tree representing the 4,366 genomes was constructed using 2,084 representative genomes due to computational limitations (Figure 2). All existing taxonomic species in the dataset were represented (see methods, Supplementary Table 2). The features of the core genome tree correlated with previously reported findings. For example, *A. hydrophila* and *Aeromonas dhakensis* formed a common clade, consistent with reclassification of *A. dhakensis* as a separate species from *A. hydrophila* subsp. *dhakensis* (51). Similarly, *Aeromonas allosaccharophila* and *A. veronii* clades are adjacent on the tree. Other relations such as the position of *Aeromonas taiwanensis* and *Aeromonas sanarellii* as neighbouring branches to *A. cavaie* were also consistent with previous findings (52).

**Figure 2.**
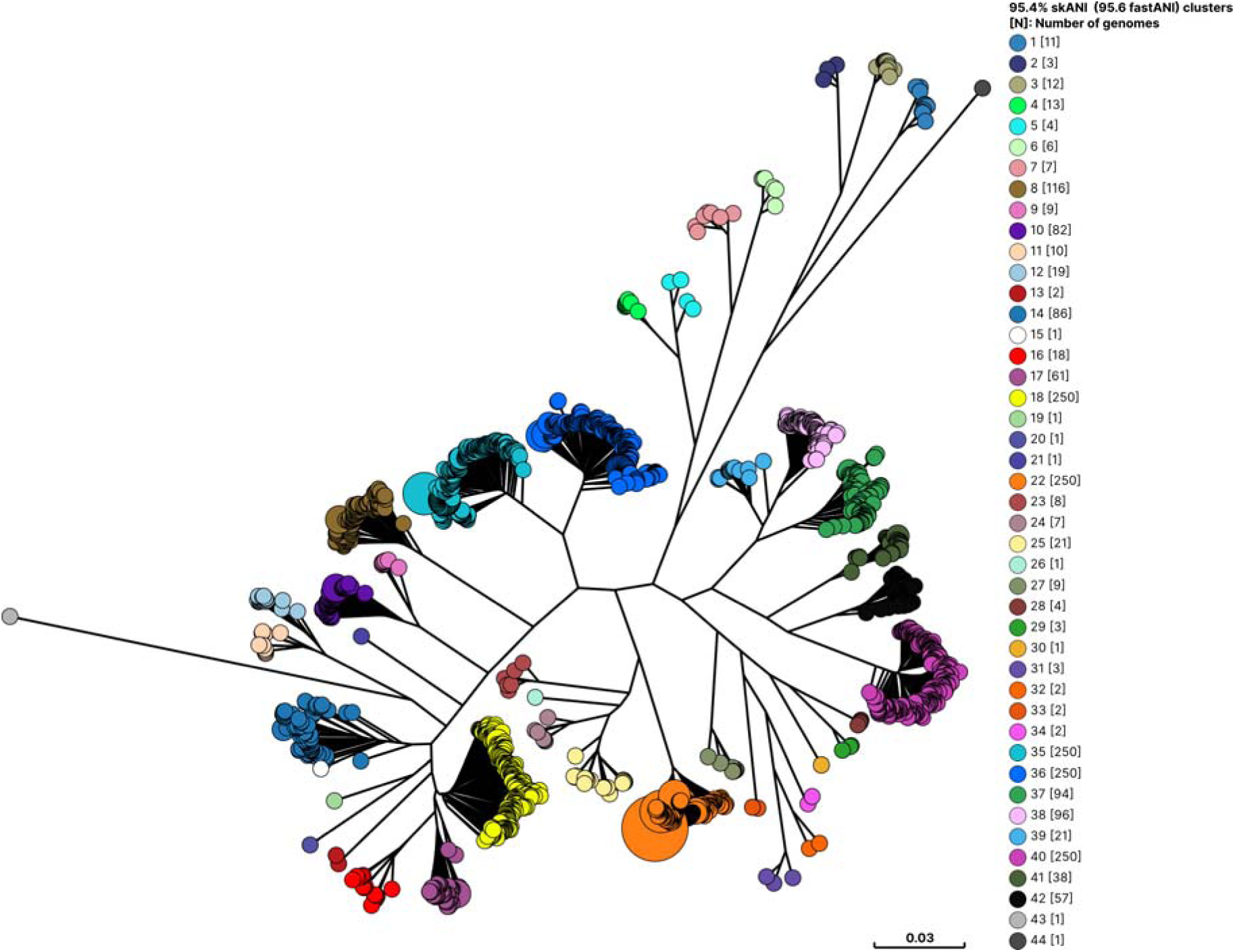
Core genome phylogenetic tree representing the 4,366-genome dataset overlaid with the 44 clusters delineated at average nucleotide identity (ANI) cutoff of 95.4% (skANI values) A total of 2,084 genomes representing the 44 clusters defined at skANI threshold 95.4% (fastANI 95.6%) were used to construct this core genome tree. The maximum likelihood tree was constructed using concatenated alignments of 673 *Aeromonas* genus core genes as input for IQ- TREE v3.0.1 and visualised with GrapeTree v1.5.0. Branch support was assessed using 1000 ultrafast bootstrap replicates. The ultrafast bootstrap values were 100 for 40 of these 44 clades. The bootstrap values were 97 for ANI cluster 16, and 81 between ANI clusters 19 and 20. The single genome ANI cluster 15, which did not separate from cluster 14 in this tree, had a bootstrap value of 81. Genome core alignment lengths were greater than 370 kb. Numbers in brackets following each cluster number indicates the number of genomes in the cluster present on the core genome phylogenetic tree. N: number of genomes.

Comparison the genus core-genome with the ANI thresholds showed that the skANI thresholds of 95.3% ANI and 95.4% were the two highest thresholds that best corresponded to species-level clades in the core genome phylogeny on the core genome phylogeny tree, although neither provided a perfect match. At 95.3% skANI, *Aeromonas lacus* (cluster 9) was merged into *Aeromonas jandaei* (cluster 10), and *Aeromonas mytilicola* was merged into *Aeromonas rivipollensis* (cluster 38). At the skANI 95.4% ANI threshold (fastANI 95.6%), *A. lacus* was delineated as a distinct and supported by the core genome phylogeny. However, a single genome ANI cluster (cluster 15) splits from *A. allosaccarophila* (cluster 14) without phylogenetic support. At no ANI threshold was *A. mytilicola* was separated from *A. rivipollensis*. However, higher ANI thresholds, including the previous recommended 96% ANI, produced additional clusters that were not supported by the core genome phylogeny (Figure 2 for 95.4% skANI threshold, Supplementary Figure 2 for 96% skANI threshold).

Based on these findings, we selected 95.4% skANI threshold for *Aeromonas genomic* species delineation. Although this threshold produced one cluster split not supported by the core-genome phylogeny tree, it retained all currently recognised *Aeromonas* species within the 4,366-genome dataset as distinct genomic species, whereas the 95.3% skANI threshold merged *A. lacus* with *A. jandaei*.

### *Aeromonas* genomic species defined using ANI

Genomic species were assigned to the ANI clusters defined at skANI threshold of 95.4% (fastANI 95.6%) that are supported by the genus core-genome phylogeny (Table 1 and Figure 3). The assigned genomic species were also compared with the species identifications made by GTDB and NCBI taxonomy (Table 1).

**Figure 3.**
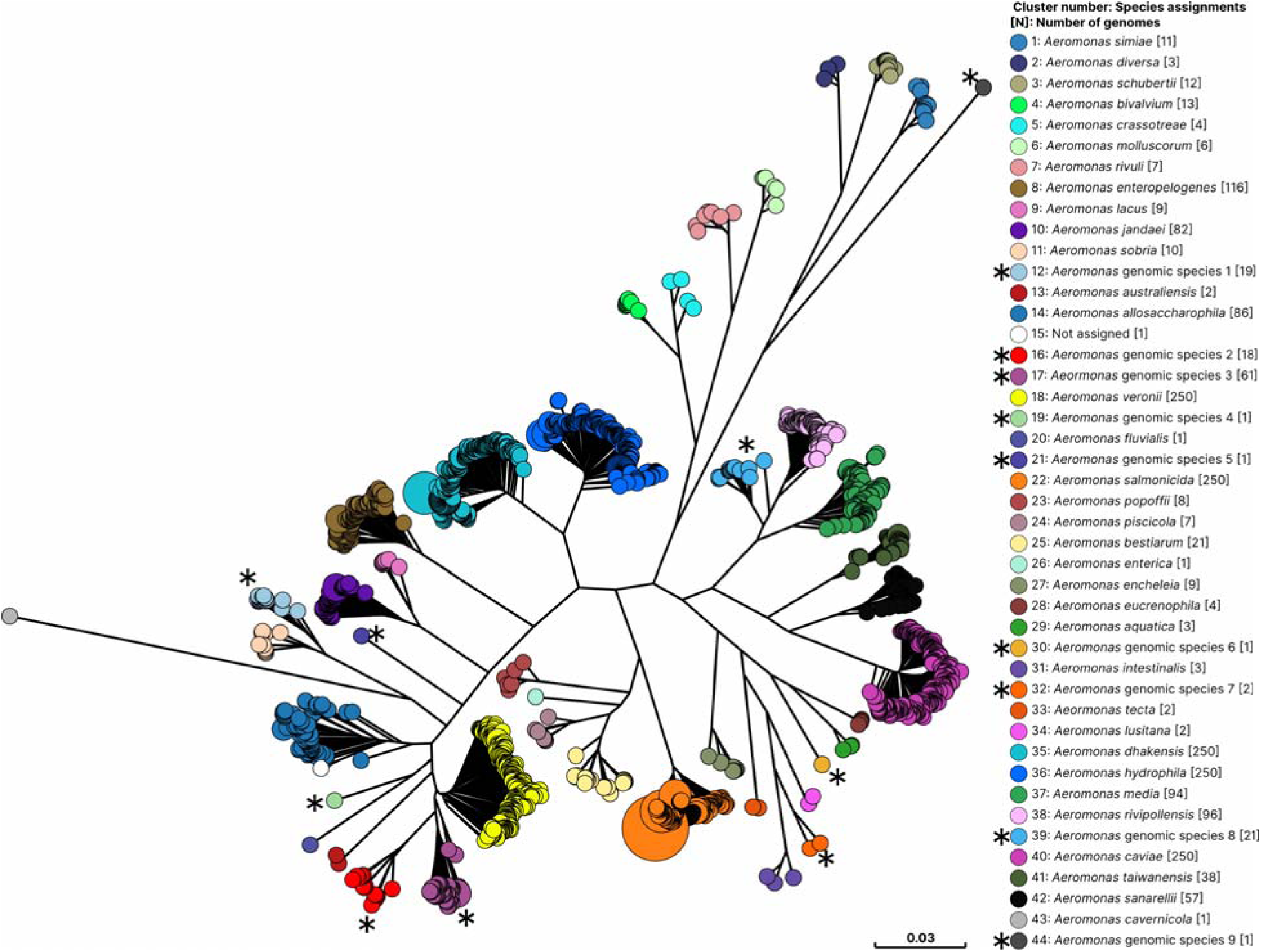
*Aeromonas* genomic species assigned using clusters delineated at the 95.4% skANI (95.6% fastANI) threshold. Phylogenetic tree presented in Figure 2 labelled with corresponding taxonomic species described in literature (Table 1). A total of 2,084 genomes representing the 44 clusters defined at skANI threshold 95.4% (fastANI 95.6%) were used to construct this core genome tree. * Denote *Aeromonas* genomic species 1-9 were assigned for the clusters with core genome phylogenetic support but no taxonomic species descriptions. *Aeromonas* genomic species 1 and 9 have descriptions as clades in species complexes (30, 55).

**Table 1.** *Aeromonas* genomic species assigned using a skANI threshold of 95.4% (fastANI threshold 95.6%) compared to species identification by NCBI and GTDB taxonomy databases.

| ANI clusters assigned in this study (95.4% skANI) | Genomic species assignment in this study (95.4% skANI) [N] | NCBI identification (96% ANI) [N] | GTDB identification (95% skANI) [N] | Reference Articles |
| --- | --- | --- | --- | --- |
| 1 | *Aeromonas simiae* [11] | *Aeromonas simiae* [11] | *Aeromonas simiae* [11] | (56) |
| 2 | *Aeromonas diversa* [3] | *Aeromonas diversa* [3] | *Aeromonas diversa* [3] | (57) |
| 3 | *Aeromonas schubertii* [12] | *Aeromonas schubertii* [11],<br>*Aeromonas* sp. [1] | *Aeromonas schubertii* [12] | (58) |
| 4 | *Aeromonas bivalvium* [13] | *Aeromonas bivalvium* [13] | *Aeromonas bivalvium* [13] | (59) |
| 5 | *Aeromonas crassostreae* [4] | *Aeromonas bivalvium* [2],<br>*Aeromonas crassostreae* [1],<br>*Aeromonas sobria* [1] | *Aeromonas sobria_B* [4] | (60) |
| 6 | *Aeromonas molluscorum* [6] | *Aeromonas molluscorum* [6] | *Aeromonas molluscorum* [6] | (61) |
| 7 | *Aeromonas rivuli* [7] | *Aeromonas rivuli* [3],<br>*Aeromonas* sp. [4] | *Aeromonas rivuli* [7] | (62) |
| 8 | *Aeromonas enteropelogenes* [116] | *Aeromonas enteropelogenes* [104],<br>*Aeromonas sobria* [1],<br>*Aeromonas* sp. [7],<br>*Aeromonas* [4] | *Aeromonas enteropelogenes* [116] | (63) |
| 9 | *Aeromonas lacus* [9] | *Aeromonas jandaei* [5],<br>*Aeromonas lacus* [2],<br>*Aeromonas* sp. [2] | *Aeromonas lacus* [9] | (28) |
| 10 | *Aeromonas jandaei* [82] | *Aeromonas jandaei* [75],<br>*Aeromonas* sp. [7] | *Aeromonas jandaei* [82] | (64) |
| 11 | *Aeromonas sobria* [10] | *Aeromonas sobria* [8],<br>*Aeromonas* sp. [2] | *Aeromonas sobria* [10] | (65) |
| 12 | *Aeromonas* genomic species 1 [19] | *Aeromonas sobria* [17],<br>*Aeromonas* sp. [2] | *Aeromonas sobria*_A [19] | (30) |
| 13 | *Aeromonas australiensis* [2] | *Aeromonas australiensis* [2] | *Aeromonas australiensis* [2] | (66) |
| 14 | *Aeromonas allosaccharophila* [86] | *Aeromonas allosaccharophila* [65],<br>*Aeromonas veronii* [11],<br>*Aeromonas hydrophila* [1],<br>*Aeromonas* sp. [8],<br>*Aeromonas* [1] | *Aeromonas allosaccharophila* [86] | (67) |
| 15 | Genomic species was not assigned | *Aeromonas allosaccharophila* [1] | *Aeromonas allosaccharophila* [1] |  |
| 16 | *Aeromonas* genomic species 2 [18] | *Aeromonas veronii* [9],<br>*Aeromonas* sp. [9], | *Aeromonas veronii*_A [18] | NA |
| 17 | *Aeromonas* genomic species 3 [61] | *Aeromonas veronii* [21],<br>*Aeromonas* sp. [40] | *Aeromonas veronii*_A [54],<br>*Aeromonas veronii* [7] | NA |
| 18 | *Aeromonas veronii* [840] | *Aeromonas veronii* [760],<br>*Aeromonas sobria* [1],<br>*Aeromonas* sp. [60],<br>*Aeromonas* [19] | *Aeromonas veronii* [840] | (68) |
| 19 | *Aeromonas* genomic species 4 [1] | *Aeromonas veronii* [1] | NA | NA |
| 20 | *Aeromonas fluvialis* [1] | *Aeromonas fluvialis* [1] | *Aeromonas fluvialis* [1] | (69) |
| 21 | *Aeromonas* genomic species 5[1] | *Aeromonas* sp. [1] | *Aeromonas* sp041929505 [1] | NA |
| 22 | *Aeromonas salmonicida* [563] | *Aeromonas salmonicida* [527],<br>*Aeromonas caviae* [1],<br>*Aeromonas eucrenophila* [1],<br>*Aeromonas* sp. [33],<br>*Aeromonas* [1] | *Aeromonas salmonicida* [563] | (70) |
| 23 | *Aeromonas popoffii* [8] | *Aeromonas popoffii* [7],<br>*Aeromonas* sp. [1] | *Aeromonas popoffii* [8] | (71) |
| 24 | *Aeromonas piscicola* [7] | *Aeromonas piscicola* [6],<br>*Aeromonas bestiarum* [1] | *Aeromonas piscicola* [7] | (72) |
| 25 | *Aeromonas bestiarum* [21] | *Aeromonas bestiarum* [16],<br>*Aeromonas hydrophila* [1],<br>*Aeromonas* sp. [4] | *Aeromonas bestiarum* [21] | (73) |
| 26 | *Aeromonas enterica* [1] | *Aeromonas enterica* [1] | NA | (60) |
| 27 | *Aeromonas encheleia* [9] | *Aeromonas encheleia* [6],<br>*Aeromonas* sp. [3] | *Aeromonas encheleia* [9] | (74) |
| 28 | *Aeromonas eucrenophila* [4] | *Aeromonas eucrenophila* [4] | *Aeromonas eucrenophila* [4] | (75) |
| 29 | *Aeromonas aquatica* [3] | *Aeromonas aquatica* [2],<br>*Aeromonas* sp. [1] | *Aeromonas aquatica* [3] | (28) |
| 30 | *Aeromonas* genomic species<br>6 [1] | *Aeromonas* sp. [1] | NA | NA |
| 31 | *Aeromonas intestinalis* [3] | *Aeromonas intestinalis* [1],<br>*Aeromonas veronii* [1],<br>*Aeromonas* sp. [1] | *Aeromonas veronii*_G [3] | (60) |
| 32 | *Aeromonas* genomic species<br>7 [2] | *Aeromonas* sp. [1],<br>*Aeromonas* [1] | *Aeromonas* sp051657805 [2] | NA |
| 33 | *Aeromonas tecta* [2] | *Aeromonas tecta* [2] | *Aeromonas tecta* [2] | (76) |
| 34 | *Aeromonas lusitana* [2] | *Aeromonas lusitana* [2] | *Aeromonas lusitana* [2] | (77) |
| 35 | *Aeromonas dhakensis* [279] | *Aeromonas dhakensis* [252],<br>*Aeromonas hydrophila* [10],<br>*Aeromonas bestiarum* [1],<br>*Aeromonas veronii* [1],<br>*Aeromonas* sp. [13],<br>*Aeromonas* [2] | *Aeromonas dhakensis* [279] | (51) |
| 36 | *Aeromonas hydrophila* [464] | *Aeromonas hydrophila* [437],<br>*Aeromonas dhakensis* [5],<br>*Aeromonas bestiarum* [2],<br>*Aeromonas enteropelogenes* [1],<br>*Aeromonas oralensis* [1],<br>*Aeromonas* sp. [18] | *Aeromonas hydrophila* [464] | (78) |
| 37 | *Aeromonas media* [94] | *Aeromonas media* [84],<br>*Aeromonas* sp. [10] | *Aeromonas media* [94] | (79) |
| 38 | *Aeromonas rivipollensis* [96] | *Aeromonas rivipollensis* [78],<br>*Aeromonas media* [9],<br>*Aeromonas mytilicola* [3],<br>*Aeromonas hydrophila* [1],<br>*Aeromonas* sp. [5] | *Aeromonas rivipollensis* [96] | (27, 80) |
| 39 | *Aeromonas* genomic species 8 [21] | *Aeromonas media* [13],<br>*Aeromonas* genomosp. *paramedia* [1],<br>*Aeromonas* sp. [6],<br>*Aeromonas* [1] | *Aeromonas* sp023093515 [21] | (31, 55) |
| 40 | *Aeromonas caviae* [1386] | *Aeromonas caviae* [1207],<br>*Aeromonas hydrophila* [4],<br>*Aeromonas media* [1],<br>*Aeromonas veronii* [1],<br>*Aeromonas* sp. [64],<br>*Aeromonas* [109] | *Aeromonas caviae* [1386] | (81) |
| 41 | *Aeromonas taiwanensis* [38] | *Aeromonas taiwanensis* [31],<br>*Aeromonas hydrophila* [4],<br>*Aeromonas* sp. [3] | *Aeromonas taiwanensis* [38] | (52) |
| 42 | *Aeromonas sanarellii* [57] | *Aeromonas sanarellii* [45],<br>*Aeromonas hydrophila* [1],<br>*Aeromonas* sp. [8],<br>*Aeromonas* [3] | *Aeromonas sanarellii* [57] | (52) |
| 43 | *Aeromonas cavernicola* [1] | *Aeromonas cavernicola* [1] | *Aeromonas cavernicola* [1] | (82) |
| 44 | *Aeromonas* genomic species<br>9 [1] | *Aeromonas* sp. [1] | *Aeromonas* sp900156095 [1] | NA |

GTDB uses a 95% ANI threshold calculated by skANI (website statement). The tool also retains recognised taxonomic species whose representative genomes may share up to approximately 97% ANI (23, 53). In contrast, NCBI taxonomy uses a 96% ANI threshold (k-mer based but unclear which tool) for prokaryotes unless there are species specific thresholds (22, 54).

A total of 44 clusters formed among the 4,366 *Aermonas* genomes at skANI threshold of 95.4% (fastANI 95.6%), of which 43 clusters were supported by the core-genome phylogeny. Thirty-four of the 43 clusters corresponded with existing taxonomic species; we assigned the name of the sole species or the predominant species in that cluster to maintain consistency and continuity with existing taxonomic species names (Table 1). The remaining nine phylogenetically supported clusters were designated as genomic species representing putative species that require further taxonomic investigation and formal classification. Cluster 15 was defined at the skANI threshold of 95.4% but not supported by the genus core-genome phylogeny. It remained as a cluster assigned to *A. allosaccharophila*, but further investigation is necessary.

Of the nine *Aeromonas* genomic species, two have existing phylogenetic descriptions in literature as species complexes. *Aeromonas* genomic species 1 (Table 1, Figure 3, ANI cluster 12), corresponds to *A. sobria* clade B from the *A. sobria* species complex (30). *Aeromonas* genomic species 8 (Table 1, Figure 3, ANI cluster 28), corresponds to *A. genomospecies paramedia* from the *A. media species complex* (31, 55). The taxonomic species *A. veronii* also splits into three species level phylogenetic clades (clusters 16-18), of which cluster 18 represented the majority of *A. veronii* genomes. The clade which contains the strain AMC-34 (cluster 17), was previously noted to be highly different from traditional *A. veronii* strains and likely a different species (12).

Discrepancies between the genomic species defined at 95.4% skANI and the species assignments in the NCBI database were observed at multiple places (Table 1). These discrepancies are expected given that NCBI applies a 96% ANI threshold for species identification, and we demonstrated in this study that the 96% threshold generated many additional clusters that divide existing taxonomic species without support from the core genome phylogeny. The genomic species defined at 95.4% skANI are largely consistent with the GTBD taxonomy, which uses a 95% skANI. Some differences in identification or naming were present in the five clusters including 16, 17, 26, 30 and 31. Among these, clusters 16 and 17 were derived from the existing *A. veronii* species and the remaining three clusters contained a single genome.

### The dDDH analysis of *Aeromonas* genomic species

dDDH has been regarded as a method to delineate prokaryotic species, with a dDDH value of 70% generally regarded as the threshold for delineating prokaryotic species. We performed pairwise dDDH comparisons between the representative medoid genomes of the genomic species delineated at the skANI threshold of 95.4% (fastANI equivalent of 95.6%). Pairwise dDDH values between all genomic species were below 70%, further supporting the genomic species delineation (Figure 4, medoid genomes details available Supplementary Table 3).

**Figure 4.**
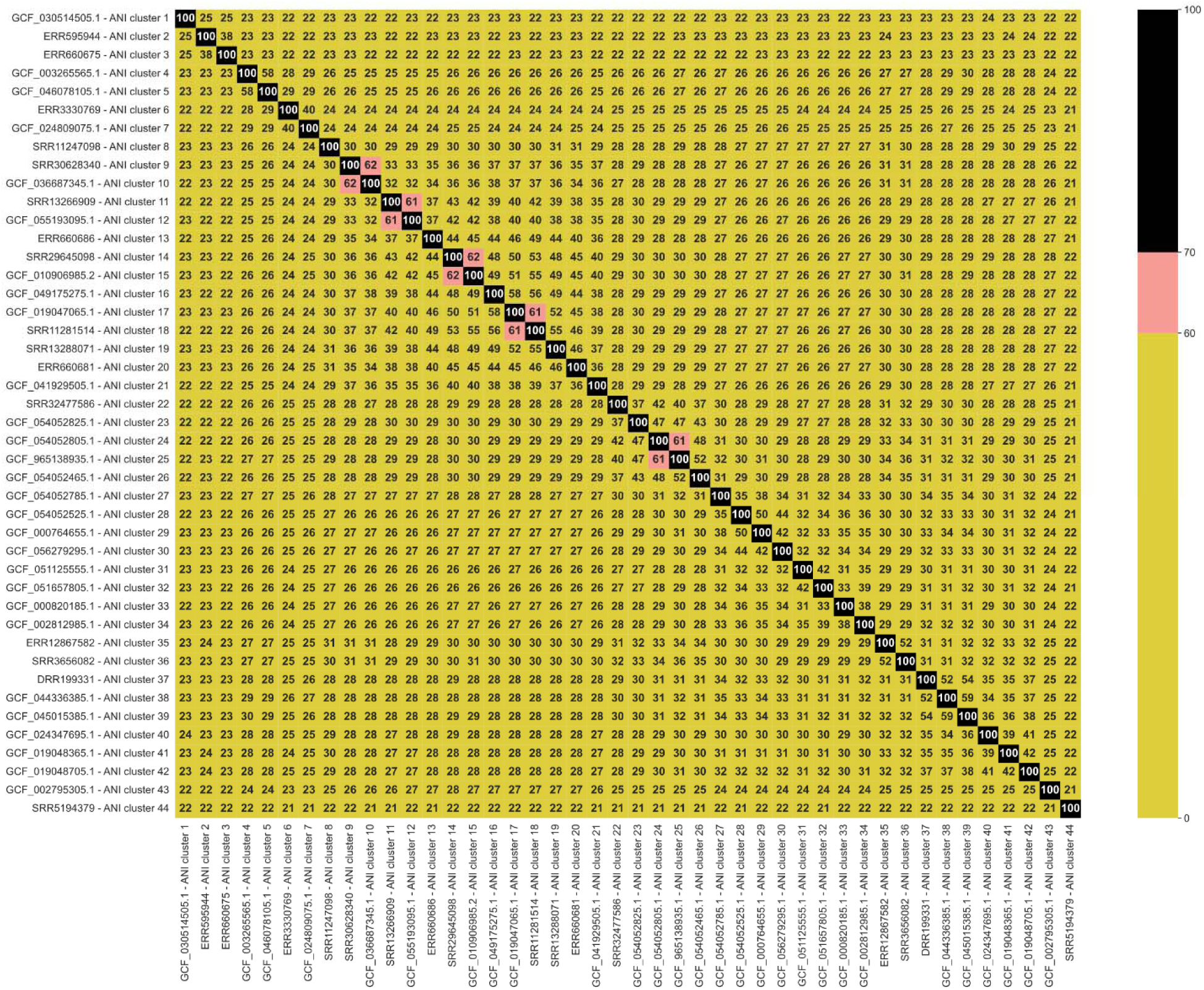
Matrix of digital DNA-DNA hybridisation (dDDH) comparisons for the 44 clusters defined at a skANI 95.4% average nucleotide identity (ANI) cutoff. The representative medoid genome of each genomic species and cluster 15 was used for pairwise comparison. Calculations of dDDH values were made using Formula of the GGDC web service and are shown as a colour coded matrix. All genomic species had interspecies dDDH value below 70%.

However, Cluster 15, a single genome ANI cluster that was not supported by the core genome phylogeny and was not defined as a genomic species, also had dDDH values less than 70% (Figure 4). This prompted us to further investigate intraspecies dDDH within *Aeromonas.* We found that seven clusters had intraspecies dDDH values below 70%. Of these clusters, five correspond to existing taxonomic species *A. jandaei, A. lacus, A. hydrophila, A. allosaccharophila, and A. veronii* (Supplementary Figure 3).

### Development of AeromonasGStyper for identification of *Aeromonas* ANI species

To facilitate the identification of genomic species for newly sequenced *Aeromonas* genomes, we developed a Python program, AeromonasGStyper. The program uses skANI or fastANI to compare a query genome with the representative medoid genome of each genomic species established in this study. Query genomes are assigned to the genomic species with the highest ANI value, provided that the ANI is ≥95.4% for skANI or ≥95.6% for fastANI. Where the assigned genomic species corresponds to a recognised taxonomic species, the recognised species name is reported. Where the assigned genomic species corresponds to an additional genomic species established in this study that does not correspond to a recognised taxonomic species, the corresponding genomic species designation is reported. Query genomes that do not meet the assignment threshold for any representative medoid genome are reported as a potentially novel genomic species with the information of the closest genomic species. Information on the representative medoid genomes is provided in Supplementary Table 3. The tool produced outputs consistent with the 4,366 genomes delineated in this study using both skANI and fastANI.

### Validation of ANI clusters using the 3,782-genome dataset

We initially collected 3,782 *Aeromonas* genomes released in the NCBI and ENA databases before January 2026. This dataset was subsequently expanded to 4,366 genomes by including additional genomes released before May 2026, which constituted the final dataset analysed in this study. The earlier dataset was used for two validation analyses. Firstly, we examined whether the relationship between skANI and fastANI clustering was consistent across different datasets. We then assessed whether the original 3,782 genomes remained assigned to the same clusters after the dataset was expanded to 4,366 genomes.

Using the 3,782-genome dataset, skANI and fastANI showed the same overall clustering behaviour observed in the 4,366-genome dataset, with fastANI thresholds consistently 0.2% higher than the corresponding skANI thresholds. Although the exact threshold pairs producing identical clusters differed slightly between the two datasets, the overall relationship between the two methods remained consistent, demonstrating that the 0.2% threshold offset was robust to dataset expansion (Supplementary Figure 4).

Comparison of the cluster assignments of the original 3,782 genomes between the two datasets showed preservation of cluster assignments across the ANI thresholds evaluated. At the selected threshold of 95.4% skANI (95.6% fastANI), cluster assignments were completely identical between the datasets, demonstrating that the selected threshold is robust to dataset expansion (Supplementary Table 4). Across the range of ANI thresholds evaluated, the same genomic clusters were recovered in both datasets. For some clusters, the ANI threshold at which they became delineated differed by only 0.1%, likely reflecting the addition of new genomes in the expended dataset (Supplementary Figure 5). Importantly, these minor shifts did not affect the selection of the operational threshold of 95.4% skANI (95.6% fastANI).

## Discussion

As taxonomic classifications of bacteria have transitioned to use of whole genome sequences, phylogeny-based assignment of species is ideal to find common ancestors and meet phylogenetic species concepts (21). However, comparative cutoff-based classifications are used for computational efficiency and quantitative values they provide, which is necessary for practical implementations (15). The 96% ANI cutoff for *Aeromonas* species delineation was proposed in 2014, based on analysis of 58 *Aeromonas* genomes (12). ANI has also been increasingly used to identify the species of prokaryotic bacteria, with the NCBI taxonomy and GTDB providing different species identifications for *Aeromonas* due to the use of 96% and 95% ANI thresholds (22, 23).

In this study, we aimed to identify an ANI threshold that is suitable for the delineation of *Aeromonas* genomic species based on multiple criteria. These criteria included consistency between ANI tools, alignment with the core genome phylogeny and dDDH, and minimal changes to existing taxonomic species. Additionally, we have compared *Aeromonas* genomic species defined in this study with existing taxonomic species and species identified in NCBI taxonomy and GTDB databases. Finally, we developed AeromonasGStyper, a tool for ANI-based *Aeromonas* genomic species identification and delineation.

Previous investigations of *Aeromonas* species used fastANI, whilst skANI is increasingly being adopted for use due to its ability to compare metagenomic assemblies, which often have completeness below 50% (17, 24). The two tools use different algorithms, which may produce different ANI values and thresholds for *Aeromonas*. We compared both tools and found that the clusters from the skANI range of 94.6-95.4% matched the fastANI range of 94.8-95.6% for the *Aeromonas* genus. In other words, fastANI thresholds were consistently 0.2% higher than the corresponding skANI thresholds. The previously recommended 96% ANI threshold for *Aeromonas* species delineation divided many recognised *Aeromonas* taxonomic species with core genome phylogenetic support. Instead, the skANI thresholds between 95.1-95.4% (fastANI 95.3-95.6%) more closely matched the existing *Aeromonas* taxonomy and warranted further comparison with the core genome phylogeny.

Core genome comparison showed that the skANI thresholds of 95.3% and 95.4% (equivalent to fastANI thresholds of 95.5% and 95.6%, respectively) were most consistent with both the core genome phylogeny and the current taxonomy. The 95.4% skANI threshold was selected for genomic species delineation in this study primarily because it retained *A. lacus* as a distinct species, whereas *A. lacus* was merged with *A. jandaei* at the 95.3% skANI threshold. The 95.4% skANI threshold preserved all currently recognised taxonomic species (34 species) among the 4,366 genomes analysed, except for *A. mytilicola*. *A. mytilicola* was merged with *A. rivipollensis* at both the 95.3% and 95.4% skANI thresholds, with no delineating cluster at higher ANI thresholds or core genome phylogenetic support. GTDB does not support the species either, despite allowing reference species up to 97% ANI. These analyses support the investigation of several potential genomic species clusters for classification as formal taxonomic species and the reassessment of the taxonomic status of *A. mytilicola*.

Conventional genome identification methods assign bacterial species by comparing a query genome with designated reference genomes (Type strain) representing recognised species. However, this approach assumes that a single reference genome adequately represents the genomic diversity of the species, which may not be appropriate for genetically diverse genera such as *Aeromonas*. In this study, we developed an *Aeromonas* genus-specific genomic species framework in which genomic species were first defined using comprehensive analyses of ANI, core genome phylogeny, dDDH and existing taxonomy, and then represented by medoid genomes selected from each genomic species. Compared with the NCBI taxonomy, which applies a universal 96% ANI threshold and consequently divides the existing *Aeromonas* species and core genome phylogeny as demonstrated in this study, our framework produced genomic species that were consistent with the core genome phylogeny while preserving nearly all recognised taxonomic species. Although GTDB species assignments were largely consistent with our genomic species delineation, several discrepancies remained because GTDB is designed to provide a universal bacterial taxonomy rather than a classification framework optimised specifically for *Aeromonas*. Instead, the framework developed in this study provides a more representative reference for genomic species assignment within the genus *Aeromonas*.

We also compared dDDH values between the representative genomes of the genomic species defined at 95.4% skANI threshold. Whilst the interspecies dDDH values were all below 70%, we also observed intraspecies dDDH values below 70% in several ANI clusters defined at 95.4% skANI threshold, five of which were established taxonomy species. Traditional DDH, which dDDH was modelled on, was shown to have a transitional range of 60-70% for species demarcation rather than having clear boundaries (20). Recently, a gradient of dDDH ranges between species was also found in the genus *Sphingobacterium* (83). Together, these observations support the use of an integrative framework that combines ANI, core-genome phylogeny and dDDH for genomic species delineation in *Aeromonas*, rather than relying on any single metric alone.

To facilitate genome-based classification of *Aeromonas*, we developed AeromonasGStyper, an ANI-based tool that implements the genomic species framework established in this study. Similar genome assignment approaches have been established for *Bacillus cereus* group and *Campylobacter* species with high accuracy (46, 84). AeromonasGStyper extends this approach to *Aeromonas* by providing rapid and standardised genomic species assignment using representative medoid genomes. As additional genomic species are established, the reference database of representative medoid genomes can be expanded, allowing future versions of AeromonasGStyper to incorporate newly recognised genomic diversity while maintaining the same assignment strategy.

In summary, through analysis of 4,366 *Aeromonas* genomes, we demonstrated that the previously recommended 96% ANI threshold overly divides recognised *Aeromonas* species, whereas 95.4% skANI (equivalent to 95.6% fastANI) provides a more appropriate operational threshold for the genomic species framework established in this study. This framework is supported by core genome phylogeny, dDDH and comparison with the existing taxonomy. Using this framework, we identified 34 *Aeromonas* genomic species corresponding to recognised taxonomic species and nine additional genomic species that do not correspond to the currently recognised taxonomic spices. Our analyses also supported further taxonomic reassessment of the distinction between *A. mytilicola* into *A. rivipollensis*. We further developed AeromonasGStyper, a rapid ANI-based tool for genomic species assignment using representative medoid genomes. Together, the genomic species framework and AeromonasGStyper provide a robust approach for genome-based classification of *Aeromonas*, improving species identification, comparative genomics and public health surveillance of *Aeromonas* infections.

## Author Contributions

Alex Chen Lu: Analysis, Validation, Writing – original draft, review and editing.

Peter Ruochen Wu: Methodology, Writing – review and editing.

Ruiting Lan: Methodology, Validation, Writing – review and editing.

Li Zhang: Conceptualisation, Funding acquisition, Supervision, Writing – review & editing.

## Acknowledgements

The authors would also like to thank the UNSW ResTech Technology Services Team for their ongoing assistance with the high-powered computing and data management systems (DOI: 10.26190/669X-A286).

## Data Summary

Data used available publicly accessible genomes from NCBI and ENA databases. Accession for genomes used are available in the supplementary data.

