## Supplementary Materials for "An integrated genomic framework for *Aeromonas* genomic species delineation using average nucleotide identify, core-genome phylogeny and digital DNA-DNA hybridization"

Supplementary Table 1. Quality checks for *Aeromonas* genomes

| **NCBI User Submitted Species** | **Genome length upper limit (bp)** | **Genome length lower limit (bp)** | **Contig limit** | **Number of contigs covering 50% of genome** | **N50 contig length (bp)** |
| --- | --- | --- | --- | --- | --- |
| *Aeromonas caviae* | 4913170.6 | 4116357.6 | 517.1 | 55.0 | 155490.5 |
| *Aeromonas veronii* | 4956681.8 | 4216459.8 | 182.8 | 22.3 | 246290.0 |
| *Aeromonas hydrophila* | 5316389.6 | 4402328.6 | 179.3 | 20.5 | 563090.0 |
| *Aeromonas dhakensis* | 5093855.3 | 4524197.3 | 126.8 | 15.5 | 691348.5 |
| *Aeromonas salmonicida* | 5197604.5 | 4380644.5 | 394.5 | 33.0 | 182916.5 |
| All other *Aeromonas* spp. criteria | 5228938.5 | 3847566.5 | 490.0 | 67.0 | 215986.0 |

For species with more than 200 NCBI user submitted genomes, species level criteria were used. For the remaining recognised NCBI species, the other *Aeromonas* spp. criteria set was used. In addition to the quality checks listed in the table, chckM contamination and completeness criteria were also used. CheckM cutoffs included removal where genome completeness was less than 95% and contamination greater than 5%. The 4,366 genomes used in this study comprised of 144 complete genomes, 86 chromosome level genomes, and 4,136 draft genomes. These genomes were filtered from a total of 7638 genomes retrieved from NCBI and ENA databases. The cutoffs for assembly statistics data were found using interquartile ranges. Chromosome level genomes are defined in NCBI as a circularised chromosome with potential gaps in additional plasmids. Two taxonomic species with literature publications, *Aeromonas finlandiensis* and *Aeromonas ichthyocola*, were not included in the analysis in this study as their representative genomes did not pass quality checks.

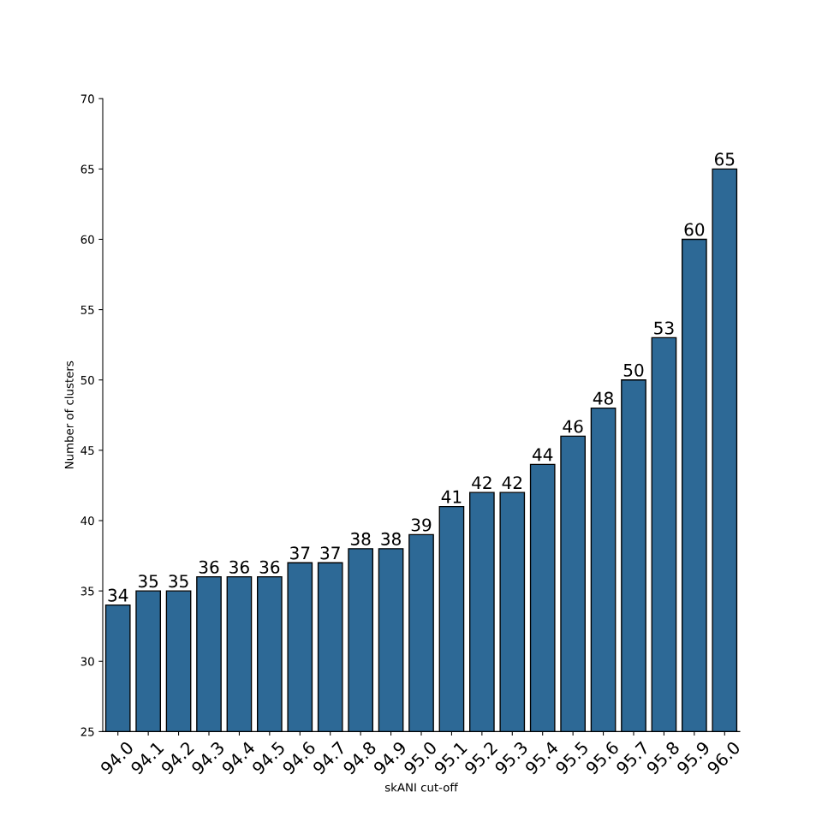

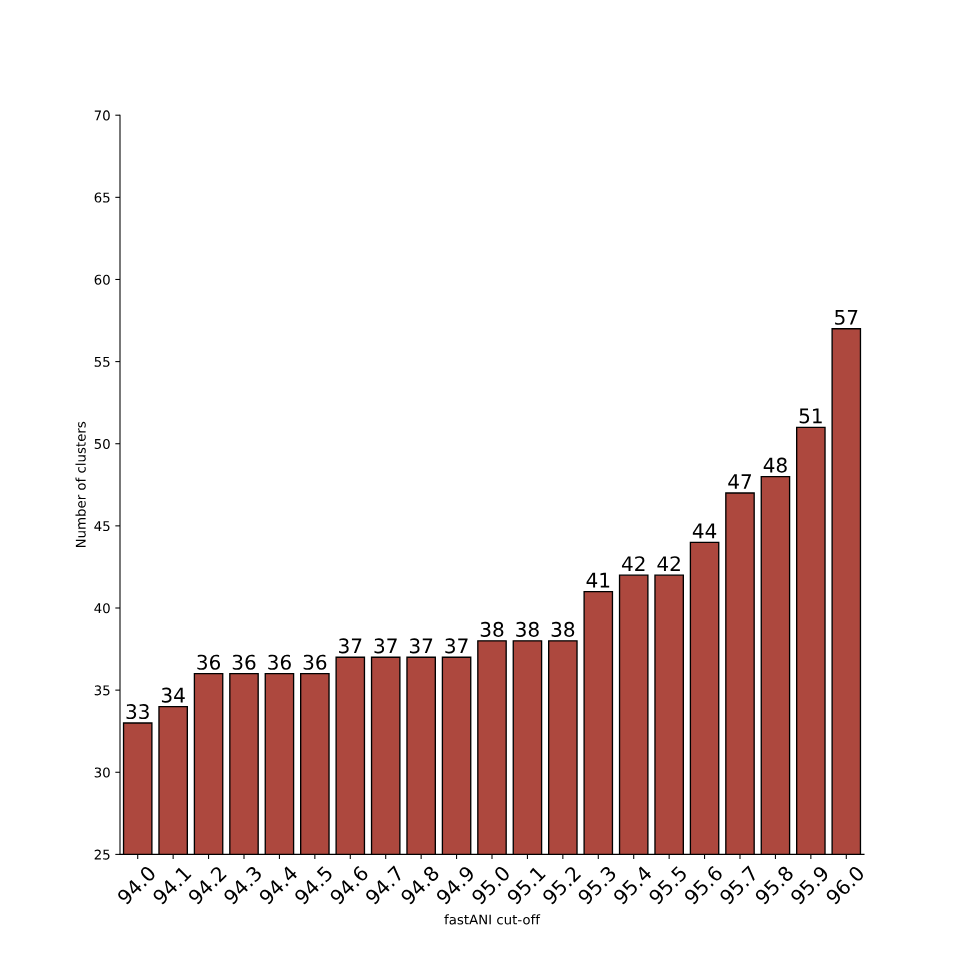

Supplementary Figure 1. Column graph representation of skANI clusters and fastANI clusters delineated at 0.1% increments

ANI pairwise comparison values generated using skANI (left, blue) and fastANI (right, red). The genomes were then delineated in clusters at 0.1% increments using average linkage clustering. These column graphs are supplementary to the Figure 1, showing cluster numbers.

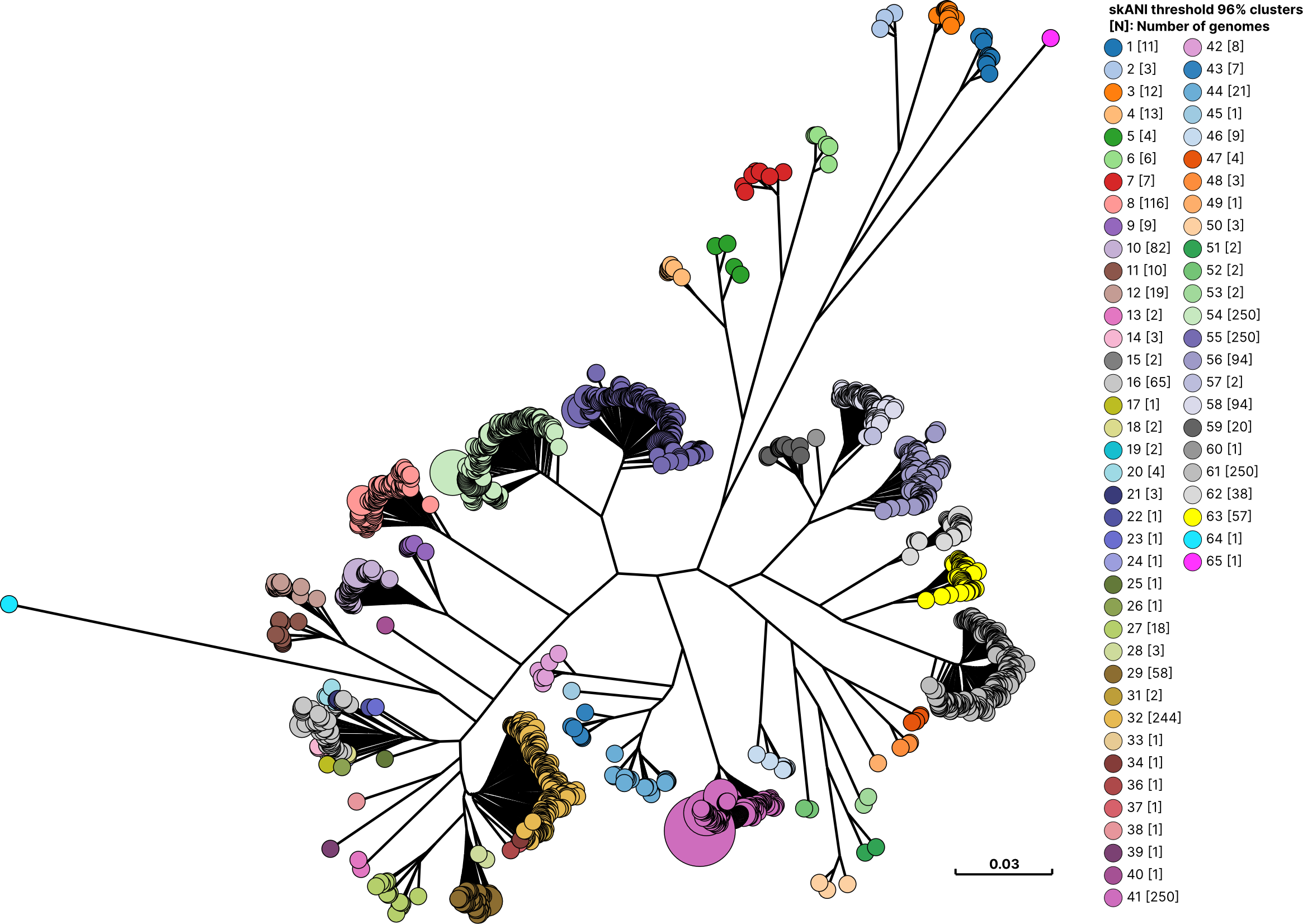

Supplementary Figure 2. Core genome phylogenetic tree representative of 4,366 genome dataset overlaid with 96% average nucleotide identity clusters (skANI)

The core genome phylogeny tree with a total of 2,083 genomes, representing the 4,366-genome dataset. The tree was overlaid with clusters at a 96% ANI cutoff using skANI. The maximum likelihood tree was constructed using concatenated alignments of 673 *Aeromonas* genus core genes as input for IQ-TREE v3.0.1 and visualized with GrapeTree v1.5.0. Not all additional clusters at 96% ANI were represented in the core genome tree. However, the additional clusters at 96% ANI do not align with the phylogenetic clades. Multiple species were separated including *A. veronii*, *A. allosaccharophila*, *A. media*, and *A. rivipollensis*,

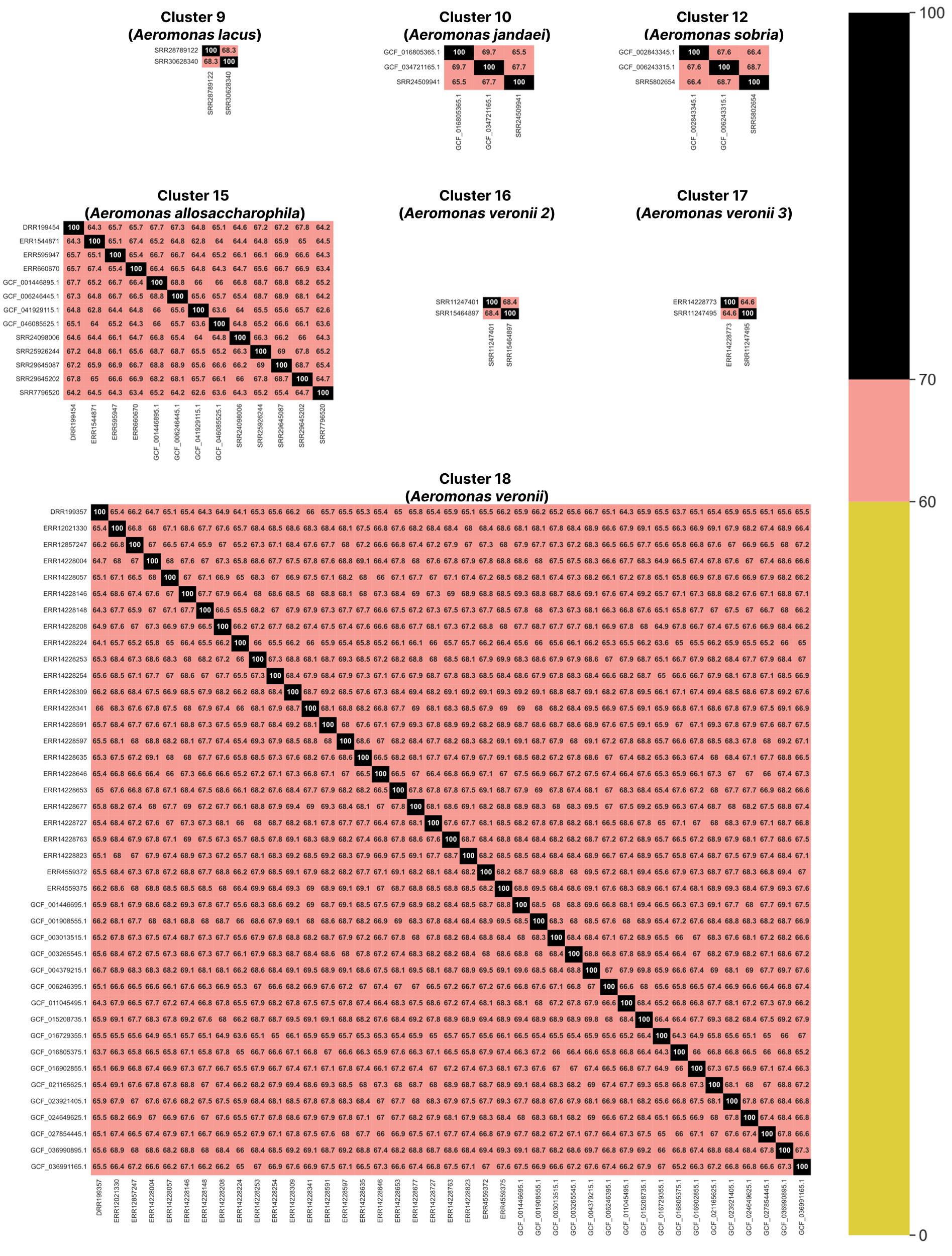

**Supplementary Figure 3. Intraspecies dDDH values below 70% observed in seven clusters delineated by skANI threshold of 95.4%**

Further dDDH comparisons were made within ANI clusters delineated at skANI threshold of 95.4% (fastANI threshold of 95.6%). It was found that seven clusters, five corresponding to existing taxonomic species, have genomes with pairwise dDDH values below 70%, but above 60%, which are presented as heatmaps.

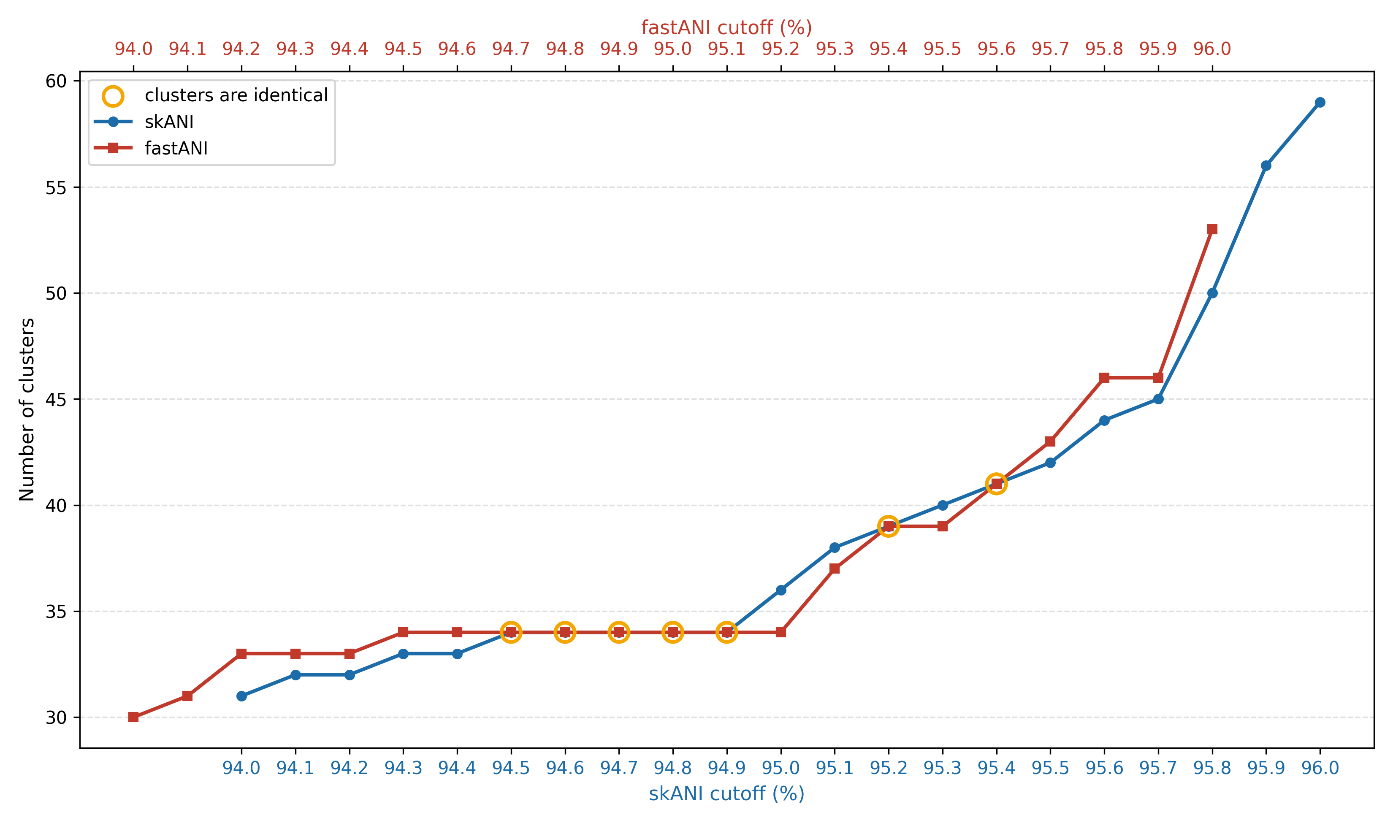

Supplementary Figure 4. Number of genome clusters delineated from the initial 3,782 *Aeromonas* genomes at different average nucleotide identity (ANI) cutoffs generated from skANI and fastANI with a 0.2% offset

ANI average linkage of 3,782 *Aeromonas* genomes using fastANI and skANI. The clusters between 94.5% to 95.4% skANI cutoffs match for most values between 94.7% to 95.6% fastANI cutoffs. The x-axis shows ANI cutoffs ranging from 94% to 96% for skANI (blue), with a 0.2% offset for fastANI (red). The y-axis shows the number of ANI clusters. Cutoffs where the number of clusters and genomes assigned to each cluster are identical between the two tools are demarcated with a yellow circle.

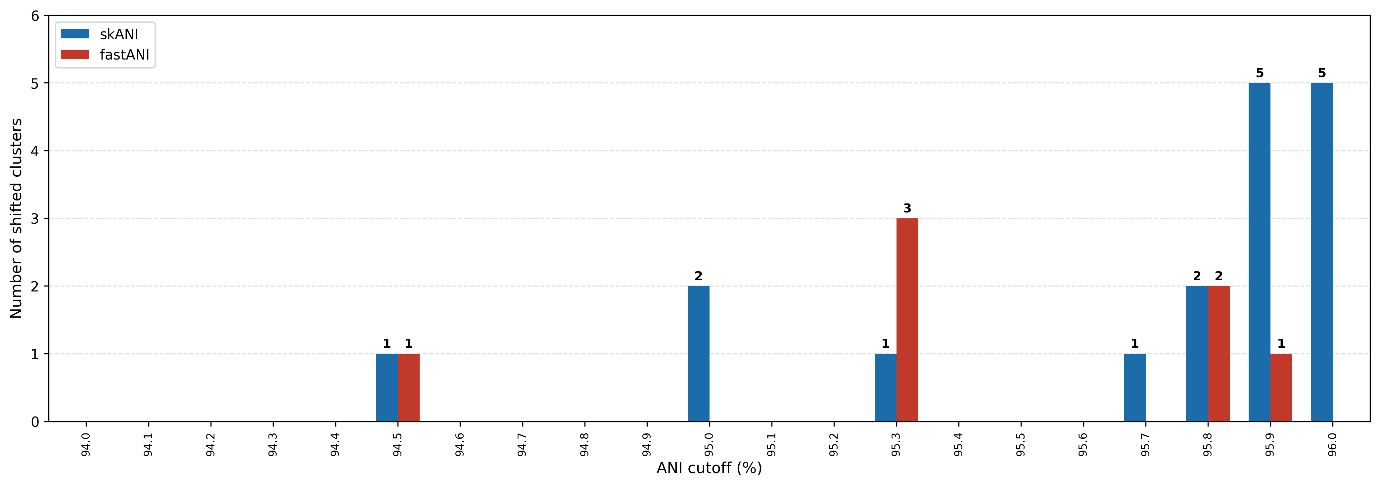

Supplementary Figure 5. Number of cluster shifts observed between the shared 3,782 genomes of the two datasets

Comparison of the cluster assignments of the 3,782 genomes between the two datasets showed that at the selected threshold of 95.4% skANI (95.6% fastANI), cluster assignments were completely identical between the datasets, demonstrating that the selected threshold is robust to dataset expansion (see Supplementary Table 2).

Across the range of ANI thresholds evaluated, the same genomic clusters were recovered in both datasets. For some clusters, the ANI threshold at which they became delineated differed by only 0.1%, likely reflecting the addition of new genomes in the expended dataset (Supplementary Figure 3). Importantly, these minor shifts did not affect the selection of the operational threshold of 95.4% skANI (95.6% fastANI).
